# ACTIVATION LOOP PHOSPHORYLATION OF A NON-RD RECEPTOR KINASE INITIATES PLANT INNATE IMMUNE SIGNALING

**DOI:** 10.1101/2021.05.01.442257

**Authors:** Kyle W. Bender, Daniel Couto, Yasuhiro Kadota, Alberto P. Macho, Jan Sklenar, Marta Bjornson, Annalise Petriello, Maria Font Farre, Benjamin Schwessinger, Vardis Ntoukakis, Lena Stransfeld, Alexandra M.E. Jones, Frank L.H. Menke, Cyril Zipfel

**Author notes:** To whom correspondence should be directed. Creoptix AG, 8820 Wädenswil, Switzerland. RIKEN Center for Sustainable Resource Science (CSRS), Plant Immunity Research Group, Yokohama, 230-0045, Japan. Shanghai Center for Plant Stress Biology, CAS Center for Excellence in Molecular Plant Sciences, Chinese Academy of Sciences, 201602, Shanghai, China. iQ Biosciences, Berkeley, 94710, California, United States (Annalise Petriello). Department of Plant Sciences, South Parks Road, University of Oxford, Oxford, OX1 3RB, United Kingdom (Maria Font Farre). Research School of Biology, The Australian National University, Acton ACT 2601, Australia. School of Life Sciences, Gibbet Hill Road, University of Warwick, Coventry, CV4 7AL, United Kingdom (Vardis Ntoukakis, Alexandra M.E. Jones).

## Abstract

Receptor kinases (RKs) play fundamental roles in extracellular sensing to regulate development and stress responses across kingdoms. In plants, leucine-rich repeat receptor kinases (LRR-RKs) function primarily as peptide receptors that regulate myriad aspects of plant development and response to external stimuli. Extensive phosphorylation of LRR-RK cytoplasmic domains is among the earliest detectable responses following ligand perception, and reciprocal transphosphorylation between a receptor and its co-receptor is thought to activate the receptor complex. Originally proposed based on characterization of the brassinosteroid receptor, the prevalence of complex activation via reciprocal transphosphorylation across the plant RK family has not been tested. Using the LRR-RK ELONGATION FACTOR TU RECEPTOR (EFR) as a model RK, we set out to understand the steps critical for activating RK complexes. While the EFR cytoplasmic domain is an active protein kinase *in vitro* and is phosphorylated in a ligand-dependent manner *in vivo*, catalytically deficient EFR variants are functional in anti-bacterial immunity. These results reveal a non-catalytic role for the EFR cytoplasmic domain in triggering immune signaling and indicate that reciprocal transphoshorylation is not a ubiquitous requirement for LRR-RK complex activation. Rather, our analysis of EFR along with a detailed survey of the literature suggests a distinction between LRR-RK complexes with RD- versus non-RD protein kinase domains. Based on newly identified phosphorylation sites that regulate the activation state of the EFR complex *in vivo*, we propose that LRR-RK complexes containing a non-RD protein kinase may be regulated by phosphorylation-dependent conformational changes of the ligand-binding receptor which could initiate signaling in a feed-forward fashion either allosterically or through driving the dissociation of negative regulators of the complex.

## Introduction

The translation of extracellular stimuli into intracellular signaling activities is carried out by myriad receptors, localized primarily at the plasma membrane. In metazoans, this role is fulfilled by proteins with diverse molecular architectures which includes ligand-perceiving G-protein coupled receptors, receptor tyrosine kinases (RTKs), Toll-like receptors, integrins, and ligand-gated ion channels. In plants, plasma membrane-localized receptor kinase (RK) complexes are the primary receivers of extracellular molecular signals, and their importance in environmental adaptation and plant development is underscored by the evolutionary expansion of RK gene families in plant genomes (Dievart et al., 2020; Dufayard et al., 2017; Hohmann et al., 2017; Shiu and Bleecker, 2003, 2001). RKs are structurally analogous to metazoan RTKs and consist of an extracellular domain that mediates ligand perception and/or protein-protein interactions, a single-pass transmembrane domain, and a cytoplasmic dual-specificity Ser/Thr and Tyr protein kinase domain (Bojar et al., 2014; Macho et al., 2015; Oh et al., 2009). Of note, plant RK cytoplasmic protein kinase domains share monophyletic ancestry with the well-known INTERLEUKIN-1 RECEPTOR-ASSOCIATED KINASES (IRAKs) that have central roles in innate immune signaling in animals (Shiu and Bleecker, 2001; Su et al., 2020) Among plant RKs, members with leucine-rich repeat (LRR) ectodomains (LRR-RKs) represent the largest sub-family and fulfill critical roles in development and stress response (Couto and Zipfel, 2016; Hohmann et al., 2017). LRR-RKs have thus been the focus of extensive biochemical and structural analyses aimed at understanding how they activate intracellular signaling in response to ligand perception (Hohmann et al., 2020; Hothorn et al., 2011; Okuda et al., 2020; Santiago et al., 2013; Sun et al., 2013b). A common mode of activation among RKs is ligand-induced heterodimerization with co-receptors. Following ligand perception, plant LRR-RKs recruit co-receptors – which are themselves LRR-RKs with short, shape-complementary ectodomains – that typically form contacts with both the ligand and the ligand-binding receptor (Hohmann et al., 2017). In this context, ligand-dependent receptor/co-receptor heterodimer formation acts as a binary switch to initiate intracellular signaling (Hohmann et al., 2020, 2018; Santiago et al., 2013; Sun et al., 2013b). Although structural analysis of receptor ectodomains has provided a detailed understanding of how receptor/co-receptor interactions occur in a ligand-dependent manner (Han et al., 2014; Hohmann et al., 2017; Hothorn et al., 2011; Santiago et al., 2013; Sun et al., 2013a, 2013b; Tang et al., 2015), much less is known mechanistically about how receptor/co-receptor dimerization activates the intracellular protein kinase activities and subsequent downstream signaling.

Early work on the brassinosteroid (BR) receptor BRASSINOSTEROID INSENSITIVE 1 (BRI1) – an LRR-RK – established that phosphorylation of both the ligand-binding receptor and co-receptor was critical for activating BR responses (Bajwa et al., 2013; Nam and Li, 2002; Wang et al., 2008, 2005). *In vitro* analysis of recombinant cytoplasmic domains revealed that BRI1 can phosphorylate its co-receptor, BRI1-ASSOCIATED RECEPTOR KINASE 1 (BAK1, also known as SOMATIC EMBRYOGENESIS RECEPTOR KINASE 3; SERK3), and that BAK1-mediated phosphorylation of BRI1 could enhance BRI1 substrate phosphorylation (Wang et al., 2008). Based on this, and the observation that both BRI1 and BAK1 are phosphorylated *in vivo* in a BR-dependent manner, ligand-triggered dimerization was proposed to facilitate reciprocal trans-phosphorylation between the receptor and co-receptor cytoplasmic domains (Chinchilla et al., 2009; Li, 2010; Wang et al., 2008), with phosphorylation events in the activation loop playing a central role. Most eukaryotic protein kinases are Arg-Asp (RD) protein kinases with Arg in the conserved subdomain VIb catalytic loop HRD motif (Hanks and Hunter, 1995) that require activation loop phosphorylation for catalytic activity (Adams, 2003; Steichen et al., 2010). Indeed, BRI1 and BAK1 are both RD protein kinases, consistent with the requirement for activation loop phosphorylation for protein function *in vivo* (Bajwa et al., 2013; Wang et al., 2019, 2008, 2005; Yun et al., 2009). However, several protein kinases, particularly in plants (Dardick and Ronald, 2006; Dardick et al., 2012), lack the conserved HRD Arg and are known as non-RD protein kinases. In both animals and plants, non-RD protein kinases have been associated with innate immune functions (Dardick and Ronald, 2006; Dardick et al., 2012). Distinct from the RD-type, non-RD protein kinases are thought not to require activation loop phosphorylation for function (Kornev et al., 2006). Although much less is known mechanistically about how the non-RD kinase are regulated, it is clear that their *in vitro* catalytic activities are low compared to their RD counterparts (Schwessinger et al., 2011). As such, it is not certain how or whether reciprocal activation loop transphosphorylation would function to activate RK complexes containing at least one non-RD protein kinase.

Targeted analysis of phosphorylation by tandem mass spectrometry of recombinant or affinity-purified proteins identified a large number of phosphorylation sites throughout LRR-RK cytoplasmic domains (Cao et al., 2013; Chen et al., 2014; Hartmann et al., 2015; Karlova et al., 2009; Mitra et al., 2015; Muleya et al., 2016; Perraki et al., 2018; Santos et al., 2009; Taylor et al., 2016; Wang et al., 2014; Xu et al., 2013, 2006; Yan et al., 2012). These targeted studies are complemented by phosphoproteomic analyses revealing multi-site phosphorylation on several RKs *in vivo* (Benschop et al., 2007; Mergner et al., 2020; Nakagami et al., 2010; Nühse et al., 2004; Sugiyama et al., 2008). By comparison to the number of phosphorylation sites documented in plant and animal systems, the vast majority of sites have not been connected experimentally to biochemical or physiological functions (Needham et al., 2019). Nevertheless, biochemical and genetic analyses have shed light on the functions of site-specific phosphorylation for some plant RKs. For example, phosphorylation of S891 in the ATP-binding loop of BRI1 inhibits its function, as indicated by increased BR responsiveness in transgenic plants expressing a non-phosphorylatable S891A mutant (Oh et al., 2015, 2012). Several LRR-RKs are phosphorylated within their intracellular juxtamembrane domains (Nühse et al., 2004), and although the specific functions of these phosphorylation events are unclear, they may control receptor stability and ligand-induced endocytic trafficking (X. Chen et al., 2010; Robatzek et al., 2006; Xu et al., 2006). In particular, phosphorylation of T705 of the rice LRR-RK XA21 inhibits immune function *in vivo* (X. Chen et al., 2010). This residue is conserved broadly across the *Arabidopsis thaliana* (hereafter, Arabidopsis) LRR-RK family, and a variant of FLAGELLIN SENSING 2 (FLS2) carrying a Thr-to-Val mutation at this position (T867V) does not undergo ligand-induced endocytosis (Robatzek et al., 2006), suggesting that phosphorylation at this site triggers receptor internalization after initiation of downstream signaling. Additional phosphorylation sites in the XA21 juxtamembrane domain are proposed to control protein stability through inhibition of cleavage by an unknown protease (Xu et al., 2006). Phosphorylation of S938 in the protein kinase domain of FLS2 positively regulates flg22 responses (Cao et al., 2013; Xu et al., 2013), but it is not clear whether this site is derived from autophosphorylation or is the target of another protein kinase *in vivo*. Evidence from analysis of the LRR-RK HAESA (HAE), which is involved in floral organ abscission, indicates that RK phosphorylation might also control substrate specificity. The HAE cytoplasmic domain is phosphorylated *in vitro* on T872 and substitution of Thr-for-Asp (T872D) specifically increases Tyr autophosphorylation activity of the protein (Taylor et al., 2016), highlighting the possibility that site-specific phosphorylation might control the dual-specificity nature of plant RKs. Although it is difficult to draw general conclusions, multiple regulatory phosphorylation sites exist on RKs, suggesting broad cellular capacity to control RK-mediated processes.

BAK1 – a common co-receptor for multiple ligand-binding LRR-RKs – is phosphorylated on multiple residues in its catalytic domain and C-terminal tail (Karlova et al., 2009; Perraki et al., 2018; Wang et al., 2008, 2014; Yan et al., 2012; Yun et al., 2009). Interestingly, a cluster of autophosphorylation sites in the BAK1 C-terminal tail (S602, T603, S604, and S612) are important for a subset of BAK1 functions, based on the conservation of a Tyr residue in subdomain VIa (which we refer to as the ‘VIa Tyr’) of the protein kinase domain of the corresponding receptor partner (Perraki et al., 2018). The BAK1 VIa Tyr (Y403) is itself phosphorylated *in vitro*, and mutation to Phe (Y403F) compromises the same subset of BAK1 functions as non-phosphorylatable mutations in the C-terminal tail cluster (Perraki et al., 2018). Although phosphorylation of Y403 or the C-tail cluster is required for full activation of immune responses, the molecular basis for their function is unknown. Intriguingly, several other RKs are phosphorylated on the subdomain VIa Tyr residue including the LysM-RK CHITIN ELICITOR RECEPTOR KINASE 1 (CERK1) and the B-type lectin S-domain RK LIPOOLIGOSACCHARIDE-SPECIFIC REDUCED ELICITATION (LORE; (Liu et al., 2018; Luo et al., 2020; Suzuki et al., 2018, 2016). For both CERK1 and LORE, phosphorylation of the VIa Tyr is required for activation of ligand-induced responses, suggesting a conserved function of this residue in RKs with diverse ectodomain architectures. The LRR-RK ELONGATION FACTOR TU RECEPTOR (EFR) is also phosphorylated on the VIa Tyr (Y836), and mutation to Phe (Y836F) abolishes ligand-dependent EFR Tyr phosphorylation and downstream signaling, suggesting that Y836 phosphorylation is required for activation of the receptor complex (Macho et al., 2014). The conservation of VIa Tyr phosphorylation on plant RKs is intriguing, although no biochemical function has yet been assigned to this important phosphorylation site.

Among the best described physiological roles for RKs is in activating cell-surface immunity where they function as pattern recognition receptors (PRRs) and perceive pathogen-associated molecular patterns (PAMPs) or host-derived damage-associated molecular patterns (DAMPs) (Couto and Zipfel, 2016; Kanyuka and Rudd, 2019). PAMP and DAMP perception sets in motion a battery of signaling events including a BOTRYTIS-INDUCED KINASE 1 (BIK1)-dependent apoplastic oxidative burst, calcium (Ca^2+^) influx (and activation of Ca^2+^-dependent protein kinases), and BIK1-independent initiation of mitogen-activated protein kinase (MAPK) cascades that collectively drive transcriptional reprogramming to ultimately halt pathogen ingress (Kadota et al., 2014; Li et al., 2014; Rao et al., 2018; Thor et al., 2020; Yu et al., 2017). In Arabidopsis, the LRR-RKs FLS2 and EFR function as PRRs to perceive the PAMPs flagellin (or the derived peptide flg22) and elongation factor thermo-unstable (EF-Tu; or the derived peptide elf18), respectively (Gómez-Gómez and Boller, 2000; Zipfel et al., 2006). Both receptors form a ligand-dependent complex with the co-receptor BAK1 or other members of the SERK subfamily (Chinchilla et al., 2007; Heese et al., 2007; Roux et al., 2011; Schulze et al., 2010). Phosphorylation of both receptor complex components occurs soon after PAMP perception and is required for downstream signaling (Macho et al., 2014; Perraki et al., 2018; Schulze et al., 2010; Schwessinger et al., 2011). EFR and FLS2 are substrates of BAK1, as is the receptor-like cytoplasmic kinase (RLCK) BIK1 (Lu et al., 2010; Schwessinger et al., 2011; Wang et al., 2014), suggesting that the majority of early activating phosphorylation events are catalyzed by BAK1 – a notion that is further supported by the dominant negative effect of catalytically inactive BAK1 mutants on PAMP signaling (Schulze et al., 2010; Schwessinger et al., 2011).

Owing to the exogenous nature of their cognate ligands, PRRs serve as a useful model to understand the biochemical mechanisms regulating receptor activity since it is possible to study acute responses to ligand perception. We previously reported on the unidirectional phosphorylation of EFR by BAK1 *in vitro* and on the critical role of EFR Tyr phosphorylation in receptor complex activation (Macho et al., 2014; Schwessinger et al., 2011). Building on these previous studies, in the present work we use EFR as a model LRR-RK and a genetic complementation approach to dissect the steps critical for phosphorylation-mediated LRR-RK complex activation. We reveal that EFR protein kinase activity is dispensable for elf18-induced immune signaling and anti-bacterial immunity and identify phosphorylation sites on purified native EFR that regulate elf18-induced receptor complex activation. Unexpectedly, we discovered EFR activation loop phosphorylation as a critical component of receptor complex activation, indicating the non-RD protein kinases might be regulated in a manner similar to enzymes of the RD type. Collectively, our data challenge the ubiquity of reciprocal transphosphorylation as a requirement for LRR-RK complex activation and support a non-catalytic role for ligand-binding receptors with non-RD intracellular protein kinase domains. We propose a mechanism where phosphorylation-dependent conformational changes of EFR would enhance co-receptor activity – either allosterically or by triggering the dissociation of negative regulators – to initiate signaling downstream of the receptor complex.

## Results

### EFR phosphorylation in the receptor complex occurs independently of its own catalytic activity

The cytoplasmic domain of EFR contains a non-RD-type protein kinase domain with Cys (C848) in place of Arg in the catalytic HRD motif, suggesting that the EFR protein kinase domain does not require activation loop phosphorylation for function (Kornev et al., 2006). Nevertheless, the recombinant EFR cytoplasmic domain (EFR^CD^) is capable of auto-phosphorylation *in vitro* following purification from *E. coli*, and similar to RD-type protein kinases, mutation of either the proton acceptor to Asn (D849N) or the catalytic loop Lys that participates in substrate coordination (K851E) (Cheng et al., 2005) compromises the protein kinase activity of EFR^CD^ (Figure 1A) (Lal et al., 2018; Schwessinger et al., 2011). We previously observed that an immunopurified EFR-BAK1 complex was catalytically active *in vitro* (Macho et al., 2014) and thus we tested whether EFR protein kinase activity was required for *in vitro* phosphorylation of the native receptor complex. Wild-type (WT) EFR or EFR^D849N^ were immunopurified from transgenic Arabidopsis seedlings expressing green fluorescent protein (GFP)-tagged EFR variants treated with mock or 100 nM elf18 for 10 minutes and the partially purified receptor complexes were then incubated with γ^32^P-ATP to assess their protein kinase activity. As in previous studies (Macho et al., 2014), EFR immunopurified from mock-treated seedlings showed minimal phosphorylation relative to the EFR-BAK1 complex purified from elf18-elicited seedlings (Figure 1B). Both BAK1 and EFR were phosphorylated in receptor complexes immunopurified from elf18-treated seedlings. Unexpectedly, the receptor complex containing EFR^D849N^ was still catalytically active, and both EFR^D849N^ and BAK1 were phosphorylated even though lower amounts of protein were immunopurified for EFR^D849N^ versus the WT (Figure 1B). This suggests that EFR catalytic activity is not required for its phosphorylation in the active receptor complex. It is likely that phosphorylation on the EFR^D849N^-containing complex is derived from BAK1 (or related SERKs), but we cannot exclude that other protein kinases in the immunoprecipitate could be responsible.

**Figure 1.**
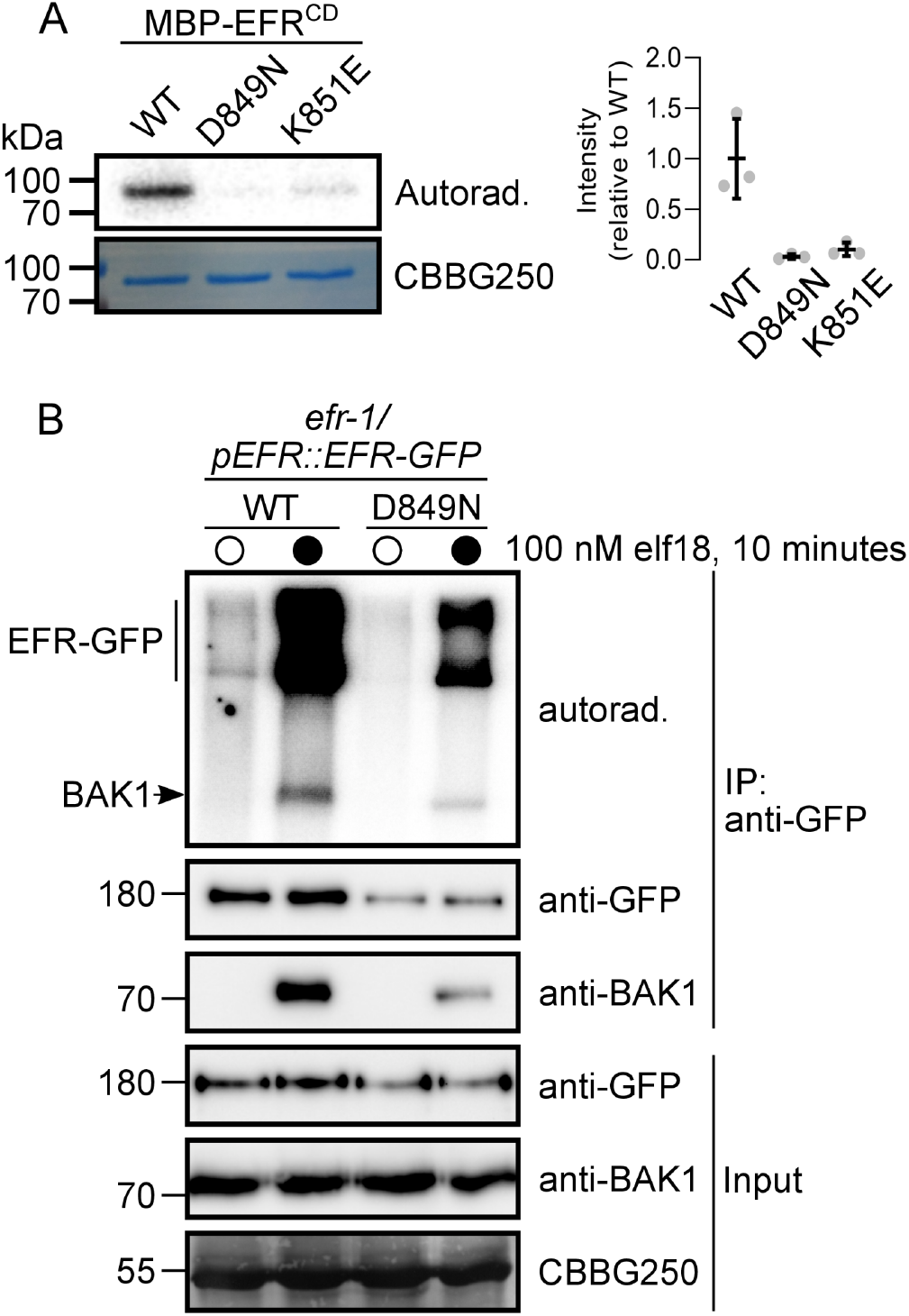
EFR is an active protein kinase but its activity is not required for phosphorylation in an isolated receptor complex. **A**, *In vitro* protein kinase activity of recombinant MBP-tagged EFR^CD^ (WT) and catalytic site mutants (D849N and K851E). Recombinant proteins were incubated with 1 μCi γ^32^P-ATP for 10 minutes and ^32^P incorporation was assessed by autoradiography. Relative quantification of ^32^P incorporation from three independent assays is shown. **B**, On-bead kinase activity assay of immunopurified EFR-GFP (mock treatment, open circles) and EFR-GFP/BAK1 (elf18-treated, closed circles) complexes purified with GFP-Trap beads. Bead-bound receptor complexes were incubated with 5 μCi γ^32^P-ATP for 30 minutes and ^32^P incorporation was assessed by autoradiography. On-bead kinase activity assays were performed three times with similar results each time.

### EFR protein kinase activity is not required for immune signaling

Because EFR protein kinase activity was not required for its phosphorylation in the isolated active receptor complex, we tested whether different catalytic site mutants of EFR could trigger the elf18-induced oxidative burst following transient expression in *Nicotiana benthamiana*, which lacks a native receptor for this PAMP. Transient expression of EFR confers perception of elf18 in *N. benthamiana* leaves as indicated by an elf18-induced oxidative burst (Zipfel et al., 2006); Figure S1). Like the WT receptor, both EFR^D849N^ and a second catalytically deficient mutant, EFR^K851E^, could activate an elf18-induced oxidative burst in *N. benthamiana* leaves but with reduced intensity or with delayed maxima compared to WT EFR (Figure S1).

We next tested whether EFR^D849N^ and EFR^K851E^ could complement the *efr-1* loss-of-function Arabidopsis mutant for elf18-induced immune signaling and anti-bacterial immunity. First, we compared WT and catalytic site mutants of EFR for activation of elf18-induced phosphorylation events by immunoblotting with phosphorylation site specific antibodies including phosphorylation of BAK1-S612, which is a marker for receptor complex formation and activation (Perraki et al., 2018), and MAPKs (Figure 2A). In transgenic Arabidopsis lines expressing EFR or the corresponding catalytic site mutants, we observed a time-dependent increase of BAK1-S612 and MAPK phosphorylation that peaked at 15 minutes following stimulation with elf18 (Figure 2A). We next measured the oxidative burst in response to elf18 treatment in the same transgenic lines. As was observed in *N. benthamiana*, both catalytic site mutants could activate an elf18-induced oxidative burst similar to the WT receptor, but with reduced intensity or with delayed maxima (Figure 2B). Notably, the total oxidative burst was reduced in transgenic plants expressing EFR^K851E^ compared to either WT or EFR^D849N^ (Figure 2B, inset); however, this difference might be attributed to reduced accumulation of the receptor in the EFR^K851E^ transgenic line (Figure 2A). Finally, we tested the effect of elf18 on seedling growth over 12 days in our complementation lines. Plants expressing the catalytically inactive variants of EFR were as sensitive to elf18 as the WT line, even at low (1 nM) concentrations of the elicitor (Figure 2C). Collectively, these experiments indicate that catalytic site mutants of EFR are competent to initiate elf18-induced signaling.

**Figure 2.**
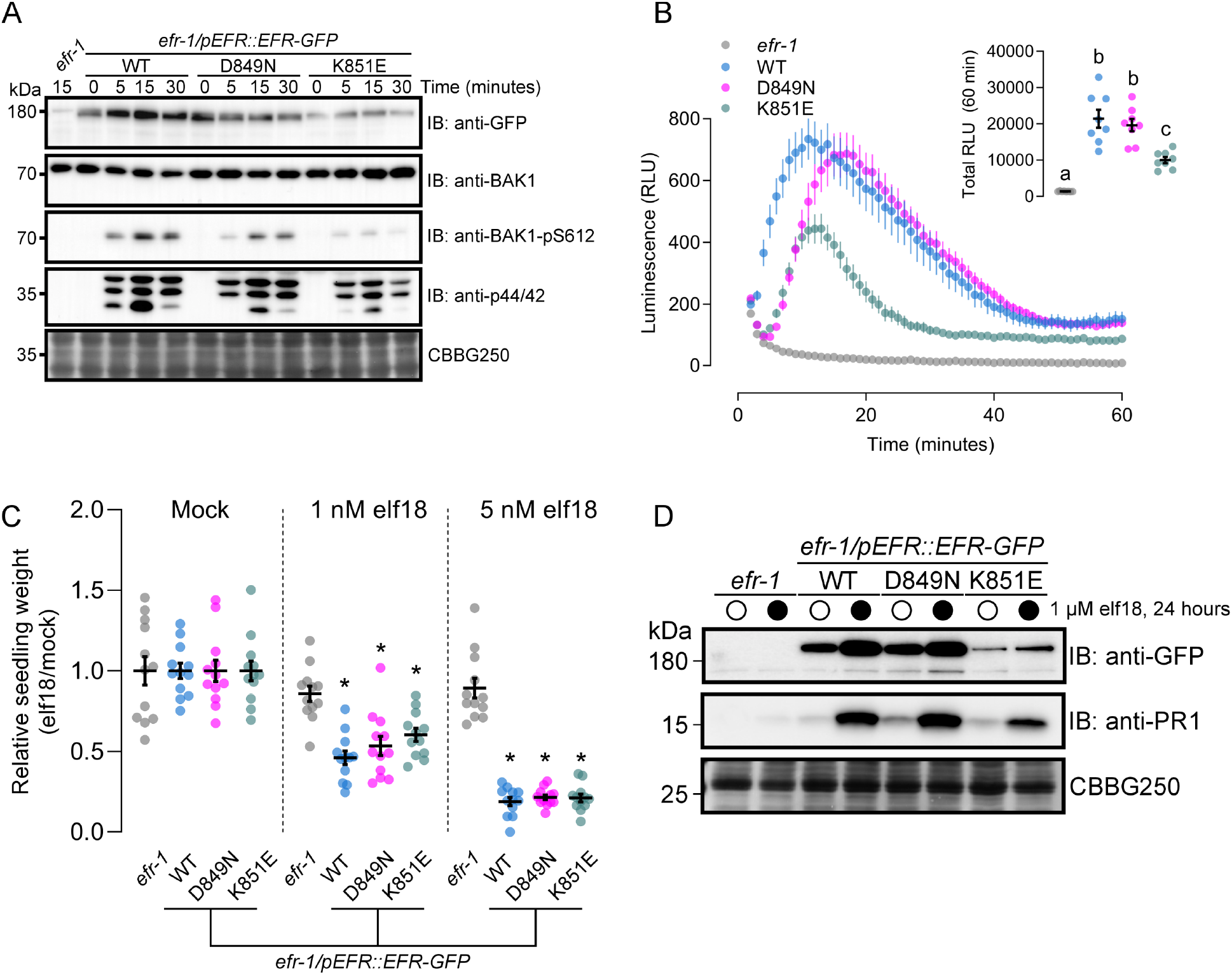
Catalytically inactive EFR variants are competent for elf18-induced PTI signaling. **A,** Immunoblot analysis of elf18-induced phosphorylation of BAK1 (anti-BAK1-pS612) and MAPKs (anti-p44/42) in 12-day-old seedlings treated with 1 μM elf18 for the indicated time. Anti-GFP shows protein accumulation of EFR and the site-directed mutants. Anti-BAK1 shows similar abundance of the co-receptor across all samples. Coomassie stain is shown as loading control (CBBG250). Blotting experiments were performed three times with similar results. **B**, Time course of the oxidative burst in leaf discs from transgenic Arabidopsis expressing EFR-GFP (WT) or kinase-dead variants (D849N or K851E) in the *efr-1* knockout background induced by treatment with 100 nM elf18. Points are mean with SEM. Inset shows mean (±SEM) of total luminescence over 60 minutes with individual data points. Different letter designations indicate statistical differences from *efr-1* (Brown-Forsythe ANOVA, n=8 leaf discs, p<0.0001, Dunnett’s multiple comparisons test). The experiment was repeated three times with similar results. **C**, Relative weight of seedlings grown in liquid media for 10 days with (1 or 5 nM) or without (Mock) the addition of elf18 peptide. Mean with SEM and individual values are shown. Asterisk indicates statistical difference from *efr-1* within a given treatment (Two-way ANOVA, n=12 seedlings, p<0.0001, Dunnett’s multiple comparison test). The experiment was repeated three times with similar results. **D**, Accumulation of PR1 protein assessed by immunoblotting with anti-PR1 antibodies 24 hours after infiltration of leaves from 3-week-old plants with mock (open circles) or 1 μM elf18 (closed circles). PR1 accumulation was assessed in three independent experiments with similar results each time.

As a second measure of long-term plant immunity signaling, we assayed salicylic acid (SA) signaling through accumulation of the SA reporter protein PATHOGENESIS-RELATED GENE 1 (PR1) (Tsuda et al., 2009; Zhang and Li, 2019) by immunoblotting with anti-PR1 antibodies. In the WT complementation line, elf18 infiltration into leaves induced robust PR1 accumulation 24 hours after treatment (Figure 3A). Like the WT, transgenic plants expressing either EFR^D849N^ or EFR^K851E^ activated PR1 accumulation in response to elf18 treatment. We additionally observed accumulation of EFR in all transgenic lines (Figure 3A), consistent with transcriptional upregulation of the receptor following elf18 perception (Bjornson et al., 2021; Zipfel et al., 2006). The accumulation of PR1 and EFR indicates that elf18-induced transcriptional responses are triggered independently of EFR protein kinase activity.

**Figure 3.**
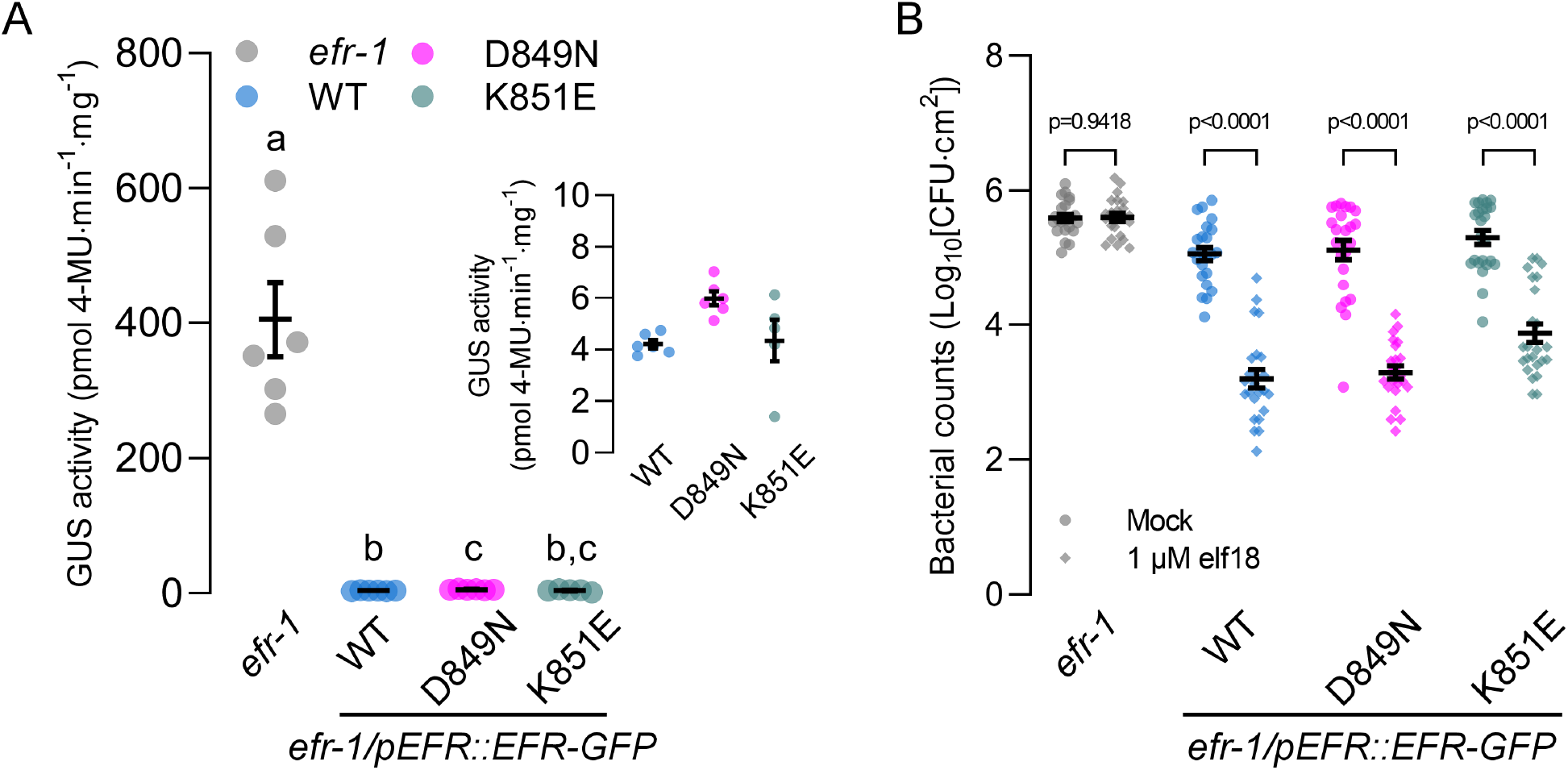
Loss of EFR kinase activity does not compromise immune responses. **A**, Fluorometric measurement of β-GUS activity in leaves of 3-week-old plants 5 days after infiltration of leaves with Agrobacterium containing the pBIN19g:GUS plasmid. Mean with SEM and individual data points are shown. Means with like letter designations are not statistically different (Brown-Forsythe ANOVA, p=0.000338, n=5 or 6 plants, Dunnett’s multiple comparisons). Inset shows zoomed y-axis to better visualize GUS activity in the complementation lines. The experiment was repeated three times with similar results. **B**, Growth of *Pto* DC3000 two days after infiltration in leaves pretreated with either mock or 1 μM elf18 for 24 hours. Mean with SEM and individual data points (n=23 or 24 plants) from three pooled independent experiments are shown. p values are derived from the comparison between elf18 pretreatment and mock, separately for each genotype as described in Experimental Procedures.

### EFR kinase activity is dispensable for anti-bacterial immunity

It is possible that although immune signaling is intact, anti-bacterial immunity could be compromised without catalytically active EFR in the receptor complex. We therefore tested whether catalytic site mutants of EFR were functional in two different pathogen infection assays: *Agrobacterium*-mediated transient transformation of Arabidopsis leaves and elf18-induced resistance to *Pseudomonas syringae* pv. *tomato* DC3000 (*Pto* DC3000) infection (Zipfel et al., 2006). In Arabidopsis, perception of EF-Tu from *Agrobacterium tumefaciens* suppresses transient transformation (Zipfel et al., 2006). To test whether EFR protein kinase activity is required to suppress transient transformation, we infiltrated leaves of *efr-1* or our complementation lines with *Agrobacterium* carrying a binary plasmid containing a β-glucuronidase (GUS) transgene (*Agrobacterium/pBIN19g:GUS*). As a proxy for transformation efficiency, we measured GUS activity in leaf extracts using a quantitative fluorometric assay (Jefferson et al., 1987). In the *efr-1* knockout line, we consistently observed GUS activity in extracts from leaves infiltrated with *Agrobacterium/pBIN19g:GUS* (Figure 3B). By comparison, GUS activity in leaf extracts from transgenic plants expressing WT EFR or the catalytic site mutants was roughly 100 times lower (Figure 3B, inset). These results indicate that catalytically deficient variants of EFR can restrict *Agrobacterium*-mediated transient transformation of Arabidopsis leaves similar to the WT receptor.

Finally, we tested whether elf18 responses triggered by the EFR catalytic site mutants could restrict *Pto* DC3000 infection. To this end, we pressure infiltrated leaves of *efr-1* and the complementation lines with either mock (sterile ddH_2_0) or 1 μM elf18, and then 24 hours later pressure infiltrated *Pto* DC3000. After two days, we measured pathogen levels by colony counting (Figure 3C). For *efr-1* knockout mutants, pathogen titer was similar in mock- and elf18-treated plants. In contrast, for the WT and both catalytic site mutant complementation lines, pre-treatment of leaves with elf18 resulted in restriction of bacterial replication compared to the mock treatment (Figure 3C), indicating that elf18-induced immune responses triggered by EFR catalytic site mutants were sufficient for induced resistance to *Pto* DC3000.

Collectively, our analysis of short-(ROS, MAPK) and long-term (seedling growth inhibition, PR1 accumulation, transient transformation, induced resistance) immune responses in transgenic plants expressing EFR^D849N^ or EFR^K851E^ demonstrate that elf18-triggered immunity does not require the catalytic activity of its cognate receptor EFR.

### Ser/Thr phosphorylation regulates EFR-mediated elf18 responses

Given that EFR is phosphorylated in the active elf18-EFR-BAK1 receptor complex, we aimed to identify the sites of phosphorylation and to test whether phosphorylation regulates elf18 responses in a site-specific manner. To identify phosphorylation sites on EFR, we immunopurified EFR-GFP from transgenic seedlings treated with mock or 100 nM elf18, and subjected the receptor complexes to an *in vitro* protein kinase assay upstream of phosphorylation site discovery by liquid chromatography-tandem mass spectrometry (LC-MS/MS). In total, we identified 12 high-confidence Ser and Thr phosphorylation sites distributed throughout the EFR cytoplasmic domain (Supplementary Table S1). Several of these sites were previously documented as either *in vitro* EFR auto-phosphorylation or BAK1 substrate phosphorylation sites (Wang et al., 2014), and several were documented as *in vivo* phosphorylation sites in a recent Arabidopsis phosphoproteome analysis (Mergner et al., 2020). Interestingly, some of the sites we identified only occurred on the receptor complex immunopurified from elf18-treated seedlings (Supplementary Table S1), suggesting that they may be involved in the regulation of EFR-mediated immune signaling.

To test if any of the identified EFR phosphorylation sites regulate elf18-triggered responses, we generated transgenic Arabidopsis plants expressing non-phosphorylatable (Ser/Thr-to-Ala) or phospho-mimic (Ser/Thr-to-Asp) mutants of EFR in the *efr-1* background and tested whether the mutants could trigger activation of MAPK cascades in response to elf18 treatment (Supplementary Figure S2A). Based on this screen, we identified two phosphosite mutants that completely lacked elf18-induced MAPK phosphorylation, namely EFR^S753D^ and EFR^S887A/S888A^. Transgenic plants expressing either the EFR^S887A^ or EFR^S888A^ single site mutant had reduced but not completely abolished MAPK activation, suggesting that phosphorylation of either residue could fulfill a putative regulatory function. In separate experiments, we tested the capacity of EFR phosphorylation site mutants to trigger BAK1-S612 phosphorylation and confirmed loss of MAPK activation for both the EFR^S753D^ and EFR^S887A/S888A^ receptor variants (Figure 4). BAK1-S612 phosphorylation could not be detected in crude extracts from transgenics expressing either EFR^S753D^ (Figure 4A) or EFR^S887A/S888A^ (Figure 4B) following elf18 treatment. By comparison, plants expressing EFR^S753A^ and EFR^S887D/S888D^ responded to elf18 similar to the WT complementation lines for both BAK1-S612 and MAPK phosphorylation (Figure 4A, B). Importantly, neither the transgenic expression of EFR^S753A^ or EFR^S887D/S888D^ led to constitutive MAPK phosphorylation, indicating that both mutant receptors still require ligand-triggered dimerization with BAK1 to activate downstream signaling.

**Figure 4.**
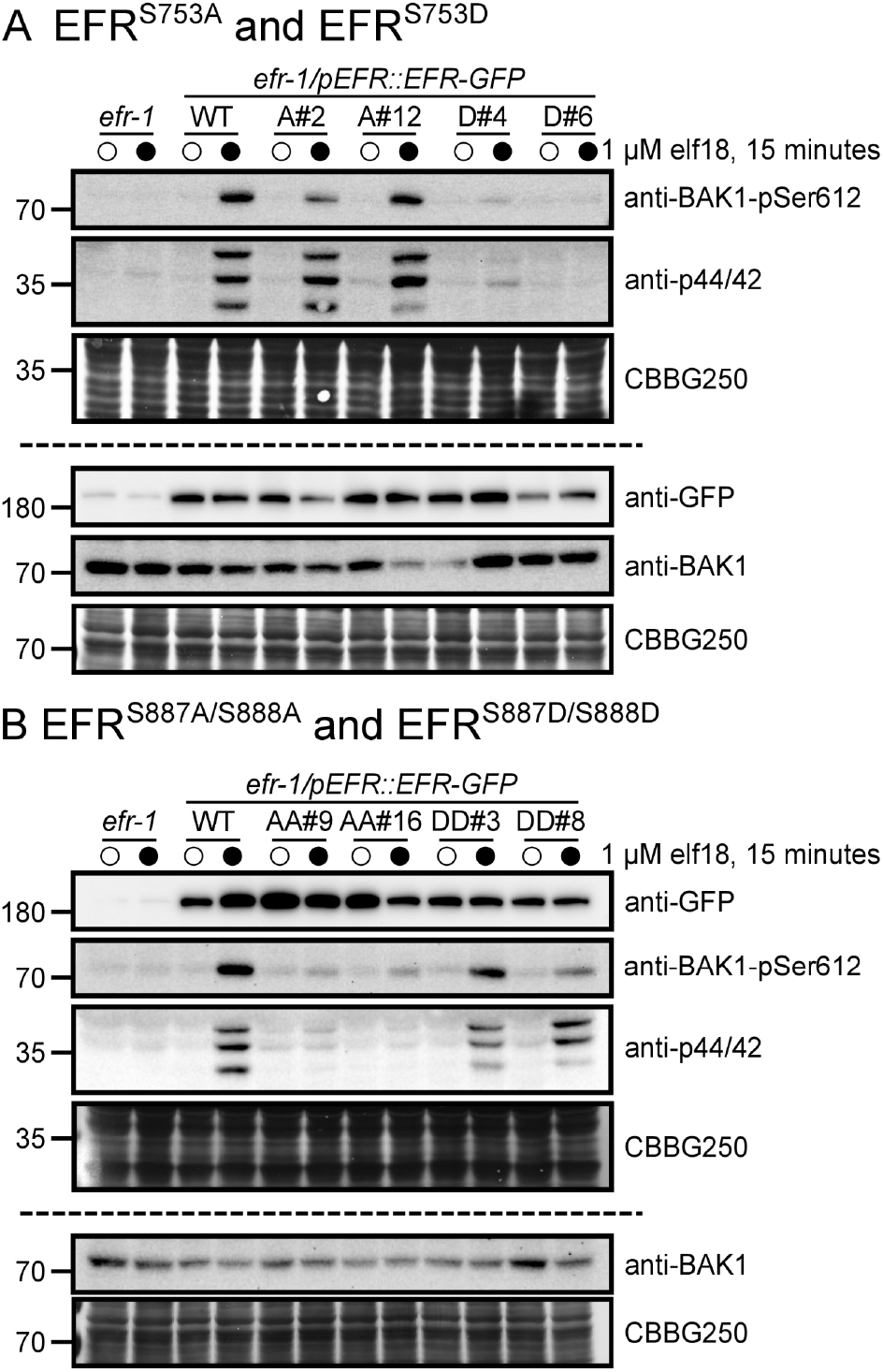
EFR phosphorylation site mutants fail to trigger ligand-induced phosphorylation events. Immunoblot analysis of elf18-induced phosphorylation of BAK1 (anti-BAK1-pS612) and MAP kinases (anti-p44/42) in 12-day-old seedlings expressing WT EFR and **A**, EFR^S753A^ (A#2, A#12) or EFR^S753D^ (D#4, D#6), or **B**, EFR^S887A/S888A^ (AA#9, AA#16) or EFR^S887D/S888D^ (DD#3, DD#8) mutants. Seedlings were treated with mock (open circles) or 1 μM elf18 (closed circles) for 15 minutes. Anti-GFP shows protein accumulation of WT EFR-GFP and the site-directed mutants. Panels above and below the dashed line represent immunoblots derived from replicate SDS-PAGE gels. Coomassie stained blots are shown as loading control (CBBG250). Experiments were repeated three times with similar results.

Next, we tested whether the EFR^S753D^ and EFR^S887A/S888A^ mutants could form functional ligand-induced receptor complexes (Figure 5). Co-immunoprecipitation experiments indicated that both EFR^S753D^ and EFR^S887A/S888A^ can form a ligand-induced complex with the co-receptor BAK1 (Figure 5A). However, by comparison to WT EFR, BAK1 co-purified with either EFR^S753D^ or EFR^S887A/S888A^ had reduced levels of S612 phosphorylation (Figure 5A), indicating that phosphorylation of S753 and S887/S888 regulate activation of the PRR complex. Additionally, we evaluated the global phosphorylation status of immuno-purified EFR or the phosphorylation site mutants by blotting with the biotinylated PhosTag reagent. We could detect elf18-inducible phosphorylation of WT EFR and the EFR^S753D^ mutant, but not EFR^S887A/S888A^ mutant (Figure 5B), suggesting a strict requirement of EFR activation loop phosphorylation for complex activation.

**Figure 5.**
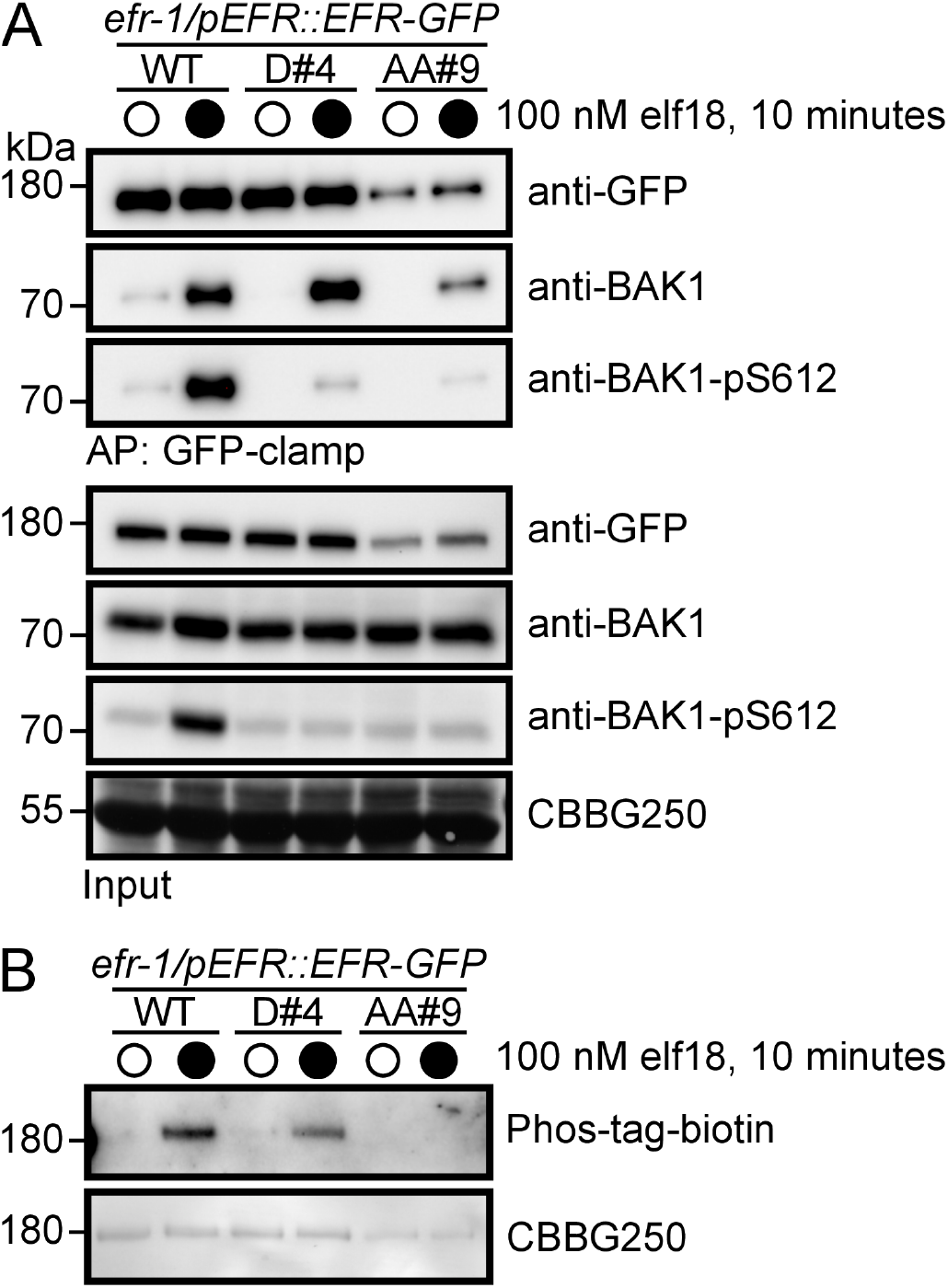
EFR phosphorylation site mutants form a ligand-induced complex with BAK1. **A**, Immunoblot analysis of elf18-induced receptor complex formation in 12-day-old seedlings expressing either WT EFR or phosphorylation site mutants (S753D, D#4; S887A/S888A, AA#9). Seedlings were treated with either mock (open circles) or 100 nM elf18 (closed circles) for 10 minutes, followed by co-immunoprecipitation with GFP-clamp beads and blotting with antibodies as indicated. **B**, Analysis of *in vivo* phosphorylation of WT EFR or phosphorylation site mutants. Seedlings were treated with either mock (open circles) or 100 nM elf18 (closed circles) for 10 minutes, followed by immunoprecipitation of GFP-tagged receptors with GFP-Trap beads. Phospho-proteins were detected using a Zn^2+^-Phos-tag::biotin-Streptavidin::HRP complex. Experiments in A and B were repeated four times with similar results.

Finally, we hypothesized that specific EFR phosphorylation sites might regulate distinct downstream pathways in a manner reminiscent of animal RTKs (Lemmon and Schlessinger, 2010). We therefore tested whether the EFR^S753D^ and EFR^S887A/S888A^ mutants were compromised in other branches of immune signalling or whether MAPK activation was the only downstream response affected. Based on our observations of receptor complex phosphorylation, we expected that other downstream responses would be similarly abolished in transgenic plants expressing either EFR^S753D^ or EFR^S887A/S888A^. Indeed, for the apoplastic oxidative burst and seedling growth inhibition, both phosphorylation site mutants were blind to elf18 treatment (Figure 6), suggesting that phosphorylation of S753 or S887/S888 does not function to regulate specific branches of immune signaling. Unlike MAPK phosphorylation, the S887D/S888D receptor variant did not fully complement *efr-1* mutants for the apoplastic oxidative burst or for seedling growth inhibition, suggesting that the double Asp mutant does not completely mimic for activation loop phosphorylation. Collectively, the loss of elf18 responses in EFR^S753D^ and EFR^S887A/S888A^ mutants indicates that the novel S753 and S887/S888 phosphorylation sites of EFR are negative and positive regulators of receptor complex activation, respectively.

**Figure 6.**
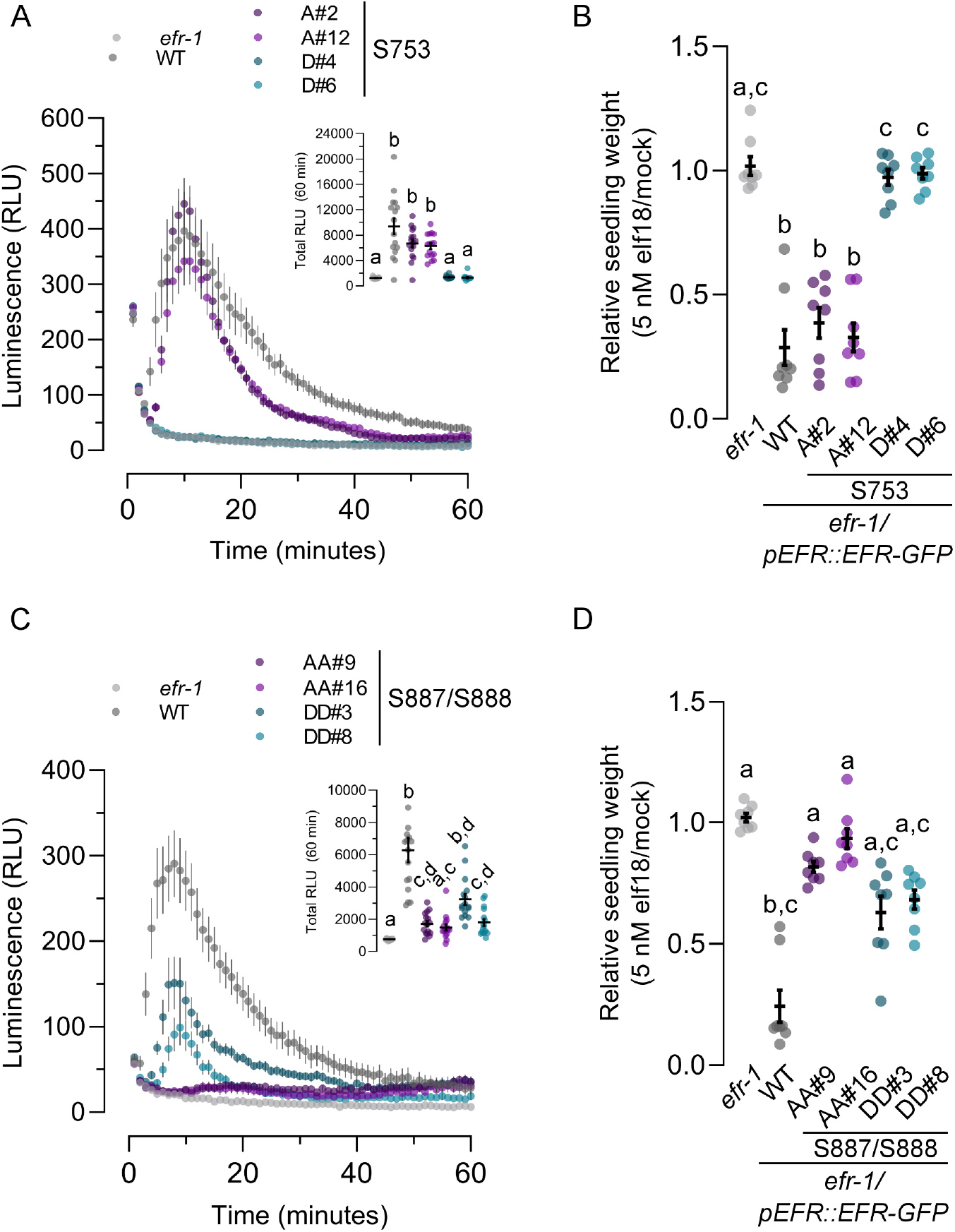
Analysis of PTI responses in EFR phosphorylation site mutants. **A**, **C**, Oxidative burst in leaf discs from the indicated genotype after treatment with 100 nM elf18. Points represent mean with SEM (n=16 leaf discs). Inset shows mean with SEM (n=16 leaf discs) of total luminescence over 60 minutes. Means with like letter designations are not statistically different (A, Kruskal-Wallis ANOVA, p<0.000001, Dunn’s multiple comparisons test; C, Kruskal-Wallis ANOVA, p<0.000001, Dunn’s multiple comparisons test). **B**, **D**, Seedling growth of the indicated genotypes in the presence of 5 nM elf18. Data are shown relative to mock treated seedlings for each genotype. Individual data points with mean and standard deviation are shown. Means with like letter designations are not statistically different (B, Kruskal-Wallis ANOVA, p=0.00001, n=8 seedlings, Dunn’s multiple comparison test; D, Kruskal-Wallis ANOVA, p<0.000001, n=8 seedlings, Dunn’s multiple comparison test). All experiments presented were repeated three times with similar results.

## Discussion

The activation of transmembrane receptors in response to exogenous and endogenous signals is a critical biochemical process in all aspects of organismal development and stress response. The plant plasma membrane is decorated with a diverse suite of RKs that perceive a wide range of ligands. The largest family of RKs in plants, the LRR-RKs, fulfill critical roles in plant development and environmental response. Members of the LRR-RK family function coordinately with co-receptors from the SERK family to activate intracellular signaling following ligand perception. While the details of ligand perception have been quantitatively described (Hohmann et al., 2017; Hothorn et al., 2011; Okuda et al., 2020; Santiago et al., 2013; Sun et al., 2013a, 2013b), much less is known about how a switch from the ligand-free apo-state to a ligand-bound activated state triggers intracellular signal transduction via the cytoplasmic protein kinase domains of the receptor and co-receptor.

In the present study, we aimed to understand the requirements for activation of LRR-RK-mediated signaling on the cytoplasmic side of the receptor complex. Using EFR as a model LRR-RK, our analyses reveal that contrary to previous reports (Lal et al., 2018; Majhi et al., 2019), catalytic activity of the ligand binding receptor is dispensable for downstream signaling. Although the consensus view is that ligand-induced dimerization triggers reciprocal transphosphorylation of receptor cytoplasmic domains, several lines of evidence suggest that transphosphorylation between the receptor and co-receptor is not required for signaling downstream of elf18 perception. First, the recombinant BAK1 cytoplasmic domain can phosphorylate the EFR cytoplasmic domain *in vitro*, but not *vice versa* (Schwessinger et al., 2011; Wang et al., 2014). Second, expression of a BAK1 kinase-inactive mutant in the null *bak1-4* background has a dominant negative effect on the elf18-induced oxidative burst (Schwessinger et al., 2011), indicating an absolute requirement for the kinase activity of BAK1 (and most likely related SERKs) for the elf18 response and suggesting that the activity of EFR is not sufficient for elf18-triggered signaling. Third, BAK1 phosphorylates the BIK1 activation loop on T237 that is required for BIK1 function (Lu et al., 2010; Xu et al., 2013), and BIK1 is the direct executor for multiple branches of immune signaling (Kadota et al., 2014; Lal et al., 2018; Li et al., 2014; Ranf et al., 2014; Thor et al., 2020). It is also noteworthy that FLS2 does not phosphorylate BIK1 *in vitro* (Xu et al., 2013), and although it has been proposed that EFR-mediated phosphorylation of BIK1 is important for immunity (Lal et al., 2018), our analysis of EFR kinase-inactive mutants indicates that this is not required for a fully functional immune response *in planta*. Collectively, these prior observations suggest that a simple phosphorylation cascade initiated by BAK1 would be sufficient to activate immunity, and that reciprocal transphosphorylation by both receptor components is not required.

Our observation that the catalytic activity of EFR is dispensable for all elf18-induced immune responses (Figures 2 and 3) argues against the ubiquity of reciprocal transphosphorylation as an activating mechanism within the plant RK family, even though formation of receptor complexes with multiple protein kinase domains is common (Couto and Zipfel, 2016). One possibility is that different activation mechanisms operate in RK complexes where both partners are RD protein kinases versus those where one partner is a non-RD protein kinase, such as the case for EFR. Although the functional significance is unknown, it is interesting that non-RD identity is broadly conserved in subfamily XII LRR-RKs that are hypothesized to function as PRRs (Dardick et al., 2012; Dufayard et al., 2017). Among reports that we could find in the published literature, with only a few notable exceptions plant RKs with RD-type intracellular protein kinase domains require their catalytic activity for function (Supplementary Table S4). By comparison, a catalytic mutant of XA21 – a non-RD PRR from rice – confers partial immunity to *Xanthomonas oryzae* pv. oryzae (F. Chen et al., 2010). FLS2 is reported to require its protein kinase activity for function (Albert et al., 2013; Asai et al., 2002; Gómez-Gómez et al., 2001; Sun et al., 2012), however, this conclusion is ambiguous since the accumulation of kinase-dead FLS2 at the protein level was not evaluated in most cases. Indeed, we previously reported that EFR expressed in *N. benthamiana* under the 35S promoter requires its kinase activity to support elf18-induced ROS, but information on expression of the catalytic mutant was lacking (Schwessinger et al., 2011). In the present work, we observe clear accumulation of both EFR^D849N^ and EFR^K851E^ associated with complementation of the *efr-1* mutant. The apparent requirement of FLS2 and EFR catalytic activity for PTI signaling reported in previous studies may thus be consequence of poor receptor accumulation under transient expression or in stable transgenic lines. Collectively this suggest that the dispensibility of catalytic function might be a common feature of non-RD protein kinases that function in immunity.

In the absence of a direct catalytic role, we foresee two possible functions for EFR in the receptor complex. First, EFR could serve as a protein-protein interaction scaffold to define specificity in activating downstream responses. In support of this, studies of chimeric receptor kinases indicate that the cytoplasmic domain of the ligand binding receptor defines signaling specificity (Hohmann et al., 2020, 2018). This suggests that the EFR cytoplasmic domain functions as a scaffold for the components required to execute immunity-specific downstream signaling. Second, besides functioning as a scaffold, the EFR cytoplasmic domain might serve to allosterically regulate BAK1 catalytic activity in the ligand-bound receptor complex. In either case, EFR phosphorylation could serve as a critical switch to activate the receptor complex and subsequent downstream events.

Even though EFR kinase activity is not required, EFR phosphorylation is critical for immune signaling (Macho et al., 2014). Here, we identified three novel regulatory phosphorylation sites on EFR, namely S753 and the S887/S888 doublet. In transgenic plants expressing either EFR^S753D^ or EFR^S887A/S888A^ we observed a loss of both BIK1-dependent (oxidative burst) and BIK1-independent (MAPK) signaling events, suggesting that the defect for both mutants occurs at the level of receptor complex activation. Interestingly, both the S753 and S887/S888 phosphorylation sites localize to subdomains that are important for regulatory conformational dynamics of protein kinases (Kornev and Taylor, 2015; Steichen et al., 2010; Taylor et al., 2015). Specifically, S753 is positioned within the αC-helix and the S887/S888 doublet within the activation loop (Supplementary Figure S2C). These residues are well conserved in Arabidopsis subfamily XIIa LRR-RKs and in PLANT ELICITOR PEPTIDE 1 RECEPTOR 1 (PEPR1), all of which are known or hypothetical PRRs (Dardick and Ronald, 2006; Dardick et al., 2012; Dufayard et al., 2017), but not in closely related subfamily XIIb members or other RD-type LRR-RKs (Supplementary Figure S2B), suggesting that these sites might be important in regulating immune signaling. Activation loop phosphorylation serves as a key regulatory switch of RD protein kinases (Moffett and Shukla, 2020; Nolen et al., 2004; Pucheta-Martínez et al., 2016; Steichen et al., 2010), and although a few non-RD protein kinases from non-plant eukaryotes are phosphorylated on their activation loops (Huang et al., 2018; Mattison et al., 2007; Waldron and Rozengurt, 2003), the functional significance of these phosphorylation events is not always well understood.

In a typical RD protein kinase, activation loop phosphorylation triggers conformational changes that establish a catalytically competent active state of the protein kinase domain (Taylor and Kornev, 2011). Based on our observation that catalytic activity is not required for EFR function, we do not think that phosphorylation of S887/S888 is required to promote EFR-mediated catalysis *per se*, but that an active-like conformation associated with activation loop phosphorylation might function in feed-forward allosteric activation, or might trigger dissociation of negative regulators of the complex (Segonzac et al., 2014). Consistent with a possible allosteric mechanism, a complex containing EFR^S887A/S888A^ is largely devoid of any phosphorylation (Figure 5), including on BAK1-S612. This suggests that phosphorylation of the EFR activation loop precedes all or most other phosphorylation on the receptor complex and that phosphorylation of EFR is required to fully activate BAK1.

Like the EFR^S887A/S888A^ non-phosphorylatable mutant, an EFR^S753D^ phospho-mimic mutant also abolished elf18 responses, but not complex formation with BAK1 (Figures 4 and 6). However, distinct from the EFR^S887A/S888A^, elf18-induced phosphorylation of EFR^S753D^ was similar to the WT (Figure 5), indicating residual protein kinase activity in the complex and phosphorylation of other sites on EFR. Interestingly, S753 is located at the N-terminal end of the ɑC-helix in the protein kinase N-lobe, a region of the protein kinase domain associated with conformational changes that mediate protein kinase activation (Taylor et al., 2015). Consistent with the requirement for EFR-BAK1 complex formation, EFR^S753A^ mutants did not display constitutive activation of any PTI responses. Although a possible mechanism to explain the impact is less clear compared to S887/S888, S753 phosphorylation could disrupt order-disorder transitions of the EFR ɑC-helix, explaining impaired activation of the EFR^S753D^-containing receptor complex. Indeed, intrinsic ɑC-helix disorder can promote an inactive state of some protein kinases, including plant RKs (Moffett et al., 2017; Shan et al., 2012), lending support to this notion.

Collectively, identification and characterization of EFR phosphorylation sites in the present work and in previous work from our lab suggests that phosphorylation-dependent conformational changes of the EFR cytoplasmic domain regulate receptor complex activation. We propose a model (Figure 7) where initial activation of the complex would occur as a consequence of EFR activation loop phosphorylation triggered by ligand-induced dimerization of EFR and BAK1. Subsequent conformational rearrangement of EFR would enhance BAK1 catalytic activity and promote VIa-Tyr phosphorylation of both complex components either allosterically, or by promoting the dissociation of components that negatively regulate BAK1. Direct phosphorylation of the ɑC-helix would fulfill an inhibitory role, and it is likely that the kinetics of S753 phosphorylation are important for this function. Importantly, our model explains the lack of requirement for the catalytic activity of the ligand-binding receptor. This alternative model for activation of LRR-RK complexes containing a non-RD protein kinase awaits further testing through a combination of time-resolved quantitative (phospho-)proteomics, homology-guided mutagenesis, and structural biology.

**Figure 7.**
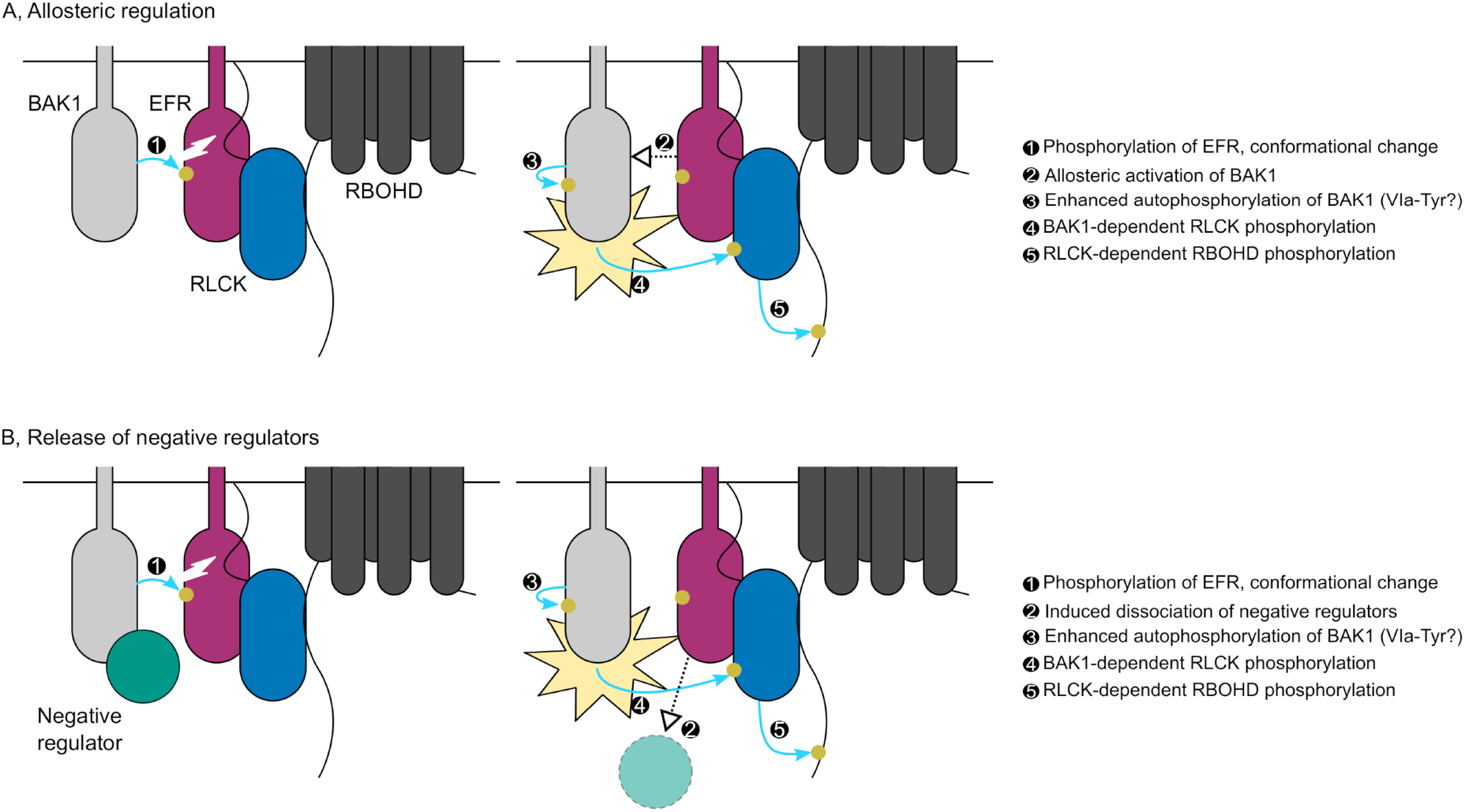
Potential mechanisms for phosphorylation-mediated activation of plant non-RD LRR-RK complexes. Ligand-triggered dimerization promotes phosphorylation of the EFR (purple) activation loop by BAK1 (light grey), inducing a conformational change of the EFR cytoplasmic domain. This conformational rearrangement feeds forward on BAK1 to enhance its catalytic activity either **A**, by direct allosteric activation of BAK1; or **B**, by triggering the release of negative regulators (teal) of BAK1 activation. Either scenario permits full phosphorylation of the complex including on the VIa-Tyr residues. After full activation, BAK1 can phosphorylate the executor RLCKs (blue) to initiate downstream signaling, for example the RBOHD (dark grey)-dependent apoplastic oxidative burst. Yellow circles and blue arrows represent simplified requirements for activation of RBOHD-dependent ROS production by phosphorylation.

## Experimental Procedures

### Plant material, growth conditions, and PAMP treatment

All genetic materials used in this study are in the Col-0 background. Complementation experiments were carried out in the *efr-1* T-DNA insertional mutant (Zipfel et al., 2006). For PAMP-induced phosphorylation (BAK1-pS612, MAPK), IP kinase, and seedling growth inhibition assays, seeds were germinated on plates containing 0.5x Murashige and Skoog (MS) basal salt mixture with 1 % (w/v) sucrose and 0.9 % (w/v) phytoagar. Growth conditions for sterile plant culture were: 120 μmol·s^−1^·m^−2^ illumination, 16 hour/8 hour day/night cycle, and a constant temperature of 22 °C. After four days of growth on agar plates, seedlings were transferred to 6-(IP kinase), 24-(PAMP-induced phosphorylation), or 48-well (seedling growth inhibition) sterile culture plates containing liquid 0.5x MS with 1 % (w/v) sucrose. For seedling growth inhibition, liquid media was supplemented with either mock (sterile ultrapure water) or elf18 peptide at the concentrations indicated in figure legends. For all experiments, seedlings were grown in liquid culture for 12 days. For PAMP-induced phosphorylation, the growth media was removed by inverting the plate on a stack of clean paper towel. Seedlings were then treated with fresh MS containing 1 μM elf18 by addition of the PAMP solution directly to the plate for the times indicated in the figures. Treated seedlings (two per treatment/time point) were dried with clean paper towel, transferred to 1.5-mL tubes, and snap-frozen in liquid nitrogen. For IP-kinase assays, seedlings from two 6-well plates (roughly 3.5 g of tissue) were transferred to 50-mL beakers containing MS and were allowed to rest for 1 hour prior to PAMP treatment. The media was then decanted and fresh MS containing mock or 100 nM elf18 was added to the beaker and was infiltrated into seedlings by the application of vacuum for 2 minutes. Seedlings were incubated in the PAMP solution for an additional 8 minutes (10 minutes treatment total) before drying with clean paper towel and snap-freezing in liquid nitrogen. All PAMP-treated plant materials were stored at −80 °C until use.

For experiments using adult (3- to 4-week-old) plants (oxidative burst, PR1 accumulation, induced resistance, transient transformation), seeds were germinated on soil and plants were grown at 22 °C/20 °C day/night temperatures with 150 μmol·s^−1^·m^−2^ illumination under a 10 hour/14 hour day/night cycle. Plants were watered automatically for 10 minutes three times per week.

### Critical reagents

Synthetic elf18 peptide was produced by SciLight Biotechnology (Beijing, China). Peptides were dissolved in sterile ddH_2_0 to a concentration of 10 mM and stored at −20 °C. Working concentrations were freshly prepared as dilutions from the stock immediately before use.

### Cloning and plant transformation

For recombinant protein expression, the EFR cytoplasmic domain was PCR subcloned from Arabidopsis cDNA using primers (Supplementary Table S2) to add KpnI and BamHI restriction sites at the 5’ and 3’ end of the amplicon, respectively. PCR products and pMAL-c4E plasmid were digested with KpnI and BamHI, digested backbone was treated with calf intestine alkaline phosphatase (CIP), and then digested PCR product and CIP-treated vector backbone were ligated with T4 DNA ligase (New England Biolabs). Ligation reactions were transformed into chemically competent *E. coli* DH10b. Individual colonies were selected for further culturing and plasmid isolation. All constructs were confirmed by DNA sequencing.

For complementation of the *efr-1* mutant with catalytically inactive and phosphorylation site variants of EFR, the EFR promoter (2.4 kb upstream of the start codon) was amplified from genomic DNA and the coding sequence from cDNA using primers for InFusion cloning (Supplementary Table S2). All constructs were confirmed by DNA sequencing prior to transformation into *Agrobacterium tumefaciens* strain GV3101. Plant transformation was carried out using the floral dip method (Clough and Bent, 1998). Transformants were selected on MS-agar plates containing 10 μg/mL phosphinothricin.

Site-directed mutagenesis to generate the catalytic site and phosphorylation site mutants was performed by rolling-circle mutagenesis using Phusion polymerase (New England Biolabs) with primers indicated in Supplementary Table S2. All mutagenized constructs were analyzed by DNA sequencing to confirm the presence of the desired mutation and the absence of off-target mutations.

### Recombinant protein expression and purification

pMAL-c4E vectors carrying in-frame fusions of the EFR cytoplasmic domain with the N-terminal maltose-binding protein (MBP) tag were transformed into Rosetta 2 cells (NEB) for recombinant protein expression. A single colony was used to inoculate a 15-mL lysogeny broth (LB) starter culture containing 100 μg/mL carbenicillin and was grown overnight at 37 °C with shaking. The next day, 1 L of LB containing 20 mM glucose and 100 μg/ml carbenicillin was inoculated with 10 mL of starter culture and was grown at 37 °C with shaking to an OD_600_ of 0.6. Recombinant protein expression was induced by the addition of 0.3 mM isopropyl β-D-1-thiogalactopyranoside (IPTG) overnight at 18 °C. Cells were pelleted by centrifugation at 5,000 rpm for 15 minutes and were then suspended in buffer containing 50 mM HEPES-NaOH pH 7.2, 100 mM NaCl, 5 %(v/v) glycerol and cOmplete EDTA-free protease inhibitor tablets (Roche).

Cells were lysed by freeze-thaw followed by sonication (four 20 second cycles with 40 second rests) and lysates were clarified by centrifugation at 35,000 x *g* for 30 minutes at 4 °C. Supernatants were adjusted to 300 mM NaCl and 2 mM DTT and were incubated with 500 μL of amylose resin (New England Biolabs) pre-equilibrated with binding buffer (50 mM HEPES-NaOH pH 7.2, 300 mM NaCl, 5 %(v/v) glycerol, 2 mM DTT) for 1 hour at 4 °C with gentle mixing. The resin was centrifuged for 10 minutes at 500 x *g* and the supernatant was discarded. The resin was suspended in 10 ml of binding buffer, mixed briefly, and centrifuged for 2 minutes at 500 x *g*. This process was repeated for a total of three washes. Bound protein was eluted from the resin by incubation for 15 minutes at 4 °C with mixing in binding buffer containing 20 mM maltose. As a final purification step, proteins eluted from amylose resin were applied to a Superdex 75 Increase size exclusion column pre-equilibrated with 50 mM HEPES-NaOH pH 7.2, 100 mM NaCl, 5 %(v/v) glycerol. Protein purity was assessed by SDS-PAGE and the concentration of peak fractions was determined by the Bradford method using bovine serum albumin as standard. Proteins samples were aliquoted and stored at −80°C until use.

### Protein extraction from plant tissues

For analysis of PAMP-induced phosphorylation by immunoblotting (MAPK and BAK1-S612), seedlings frozen in 1.5-mL tubes were pulverized with a nitrogen-cooled plastic micropestle. One hundred microliters per seedling (200 μL total) of extraction buffer containing 50 mM Tris-HCl pH 7.5, 150 mM NaCl, 2 mM EDTA, 10 %(v/v) glycerol, 2 mM DTT, 1 %(v/v) Igepal, and protease and phosphatase inhibitors (equivalent to Sigma-Aldrich plant protease inhibitor cocktail and phosphatase inhibitor cocktails #2 and #3) was added to each tube and the tissue was ground at 2000 rpm using an overhead mixer fitted with a plastic micropestle. The tubes were centrifuged at 15,000 x *g* for 20 minutes at 4 °C in a refrigerated microcentrifuge. After centrifugation, 150 μL of extract was transferred to a fresh 1.5-mL tube. Protein sample concentrations were normalized using a Bradford assay. Samples were prepared for SDS-PAGE by heating at 80 °C for 10 minutes in the presence of 1X Laemmli loading buffer and 100 mM DTT.

For co-immunoprecipitation and IP kinase assays, approximately 3.5 g of frozen tissue was ground to a fine powder under liquid nitrogen in a nitrogen-cooled mortar and pestle and then further ground with sand in extraction buffer (as described above) at a ratio of 4 mL of buffer per gram of tissue. Extracts were filtered through two layers of Miracloth and centrifuged at 25,000 x g for 30 minutes at 4 °C to generate a clarified extract.

### In vitro *protein kinase assays*

To assess the activity of recombinant MBP-EFRCD, 500 ng of purified protein was incubated in a 20-μL reaction with 1 μCi of γ^32^P-ATP, 2.5 mM each MgCl_2_ and MnCl_2_, and 10 μM ATP in 50 mM HEPES-NaOH pH 7.2, 100 mM NaCl, and 5 %(v/v) glycerol for 10 minutes at 30 °C. Reactions were stopped by the addition of Laemmli SDS-PAGE loading buffer and heating at 80 °C for 5 minutes. Reactions were separated by SDS-PAGE followed by transfer to PVDF, and exposure of storage phosphor-screen for 30 minutes. Exposed screens were imaged using an Amersham Typhoon (GE Lifesciences). Image analysis for relative quantification of ^32^P incorporation was carried out using the ImageQuant software package, with local averaging for background subtraction.

### SDS-PAGE, Immunoblotting, and chemiluminescence imaging

Proteins were separated in either 10 %(v/v) (MAPK phosphorylation), 8 %(v/v) (CoIP), or 15 %(v/v) (PR1 accumulation) polyacrylamide gels at 120 V for 95 minutes. Proteins were transferred to PVDF membranes at 100 V for 90 minutes at 4 °C followed by blocking for 2 hours at room temperature or overnight at 4 °C in 5 %(w/v) milk in Tris-buffered saline (50 mM Tris-HCl pH 7.4, 150 mM NaCl; TBS) containing 0.1 %(v/v) Tween-20 (TBS-T). Blots were probed in primary antibody according to the conditions in Supplementary Table S3, followed by washing 4 times for 10 minutes each in TBS-T. When required, blots were then probed in a 1:10,000 dilution of goat-anti-rabbit-HRP conjugate for 30 minutes to 1 hour, followed by washing 3 times for 5 minutes each in TBS-T. Blots were then washed for 5 minutes in TBS and treated with either standard ECL substrate or SuperSignal West Femto high sensitivity substrate (ThermoFisher Scientific). Blots were imaged using a Bio-Rad ChemiDoc Imaging System (Bio-Rad Laboratories). All raw images were saved in the Bio-Rad .scn format and blots were exported as 600 dpi TIFFs for preparation of figures.

For the experiments presented in Figure 2A and Figure 5A, immunoblots probed with anti-BAK1 pS612 antibodies were stripped by incubation in stripping buffer containing 214 mM glycine pH 2.2, 0.1 %(w/v) SDS, 1 %(v/v) Tween-20 4 times for 10 minutes each followed by washing in 1xTBS-T 4 times for 5 minutes each. Stripped blots were blocked overnight at 4 °C in 5 %(w/v) milk in TBS-T before probing with anti-BAK1 antibodies (Supplementary Table S3).

### Immune assays

For MAPK assays, seedlings were grown in 24-well plates (one seedling per well) as described above. Growth media was removed by inverting plates on paper towels and individual seedlings were treated as indicated in the figure legends. Two seedlings were pooled for each treatment/time point. Total proteins were extracted as described above, normalized by Bradford assay and analyzed by SDS-PAGE and immunoblotting with anti-p44/42 antibodies (Supplementary Table S3).

For analysis of the PAMP-induced oxidative burst, leaf discs from 3- to 4-week-old plants were collected into white 96-well plates using a 4 mm biopsy punch and were allowed to rest in sterile ultrapure water overnight. The next day, the water was removed and replaced with a solution containing 100 nM elf18, 1 mM luminol, and 10 μg/mL HRP (in sterile ultrapure water). Luminescence was collected for 70 minutes using a Photek system equipped with a photon counting camera.

For seedling growth inhibition assays, 4-day-old seedlings were transferred to 48-well plates (one seedling per well) containing MS with mock (sterile ddH_2_0) or elf18 at the concentration indicated in figure captions. Seedlings were grown for 10 days in the treatment solution and the weights of individual seedlings were recorded using an analytical balance.

PR1 accumulation was evaluated by immunoblotting protein extracts from leaves treated with elf18. Three leaves from 3- to 4-week-old plants were pressure infiltrated with either mock (sterile ultrapure water) or 1 μM elf18. After 24 hours, leaves were removed by cutting with sharp scissors and were snap-frozen in liquid nitrogen in 1.5-mL tubes and were then pulverized with a nitrogen-cooled plastic micropestle. Total proteins were extracted by grinding at 2,000 rpm in extraction buffer (see above) using a micropestle fixed to a rotary mixer. Protein extracts were normalized by Bradford assay and were analyzed by SDS-PAGE and immunoblotting using anti-PR1 antibodies (Supplementary Table S3).

### Agrobacterium-mediated transient transformation and induced resistance assays

Analysis of GUS activity following transient transformation of Arabidopsis leaves was performed as previously described (Jefferson et al., 1987; Zipfel et al., 2006). Briefly, *Agrobacterium tumefaciens* GV3101 carrying the pBIN19g:GUS (containing a potato intron) plasmid was infiltrated into the leaves of 3- to 4-week-old plants at an OD_600_ of 0.4. After 5 days, infiltrated leaves were removed by cutting with sharp scissors and were snap-frozen in liquid nitrogen in 1.5-mL tubes. Total proteins were extracted in GUS assay buffer (Jefferson et al., 1987) and GUS activity was measured after 30 minutes of incubation in the presence of 1 mM 4-methylumbelliferyl-β-D-glucuronide (MUG, Sigma Aldrich). Reactions were stopped by the addition of four volumes of 0.2 M Na_2_CO_3_ and fluorescence was measured in a Biotek Synergy microplate reader with excitation and emission wavelengths of 365 nm and 455 nm, respectively. The amount of 4-methylumbelliferone (4-MU) produced was measured against a standard curve of 4-MU prepared in methanol.

Induced resistance assays were performed as described previously (Zipfel et al., 2004). Briefly, 3 leaves each of 5-week-old plants grown on soil were infiltrated with a solution of 1 μM elf18 or mock (sterile ddH_2_0) in the morning. The following morning treated leaves were re-infiltrated with a suspension of approximately 10^8^ *Pseudomonas syringae* pv. tomato DC3000 (*Pto* DC3000) per mL (OD_600_=0.0002). Plants were left uncovered for two days, after which two leaf discs were harvested per treated leaf and six leaf discs pooled per plant. Colony forming units (CFU) per cm^2^ were counted through serial dilution, and statistics performed on log_10_(CFU/cm^2^) in R (R Foundation for Statistical Computing, 2020). ANOVA revealed significant effects of genotype and treatment, as well as a significant interaction (p<2.2×10-16). The effect of treatment within each genotype was estimated through estimated marginal means (package emmeans) with no correction for multiple testing (Lenth, 2020).

### Co-immunoprecipitation and IP kinase assays

Protein extracts containing GFP-tagged EFR or site-directed mutants were incubated with 20 μL of GFP-Trap beads (Chromotek) or GFP-clamp beads (Hansen et al., 2017) as indicated in figure captions for 2 hours with gentle mixing at 4 °C to immuno-precipitate receptor complexes. The beads were sedimented by centrifugation at 1000 x *g* for 5 minutes at 4 °C and were subsequently suspended in 1 mL of extraction buffer (see above). The beads were sedimented at 1,000 x *g* for 1 minute and suspended in 1 mL of extraction buffer three more times for a total of four washes. After the last wash was removed, beads were suspended in 2X Laemmli SDS-PAGE loading buffer followed by heating at 80 °C for 10 minutes. For IP-kinase assays, beads were equilibrated in 1 mL kinase assay buffer containing 50 mM HEPES-NaOH pH 7.2, 100 mM NaCl, 5 mM each MgCl_2_ and MnCl_2_, and 5 %(v/v) glycerol. The total volume of kinase assay buffer and beads was split in two and half was immediately prepared for SDS-PAGE by removal of the kinase assay buffer and heating of the beads at 80 °C for 10 minutes in 2X Laemmli SDS-PAGE loading buffer. The second half was used for an *in vitro* on-bead kinase assay.

After removal of the equilibration buffer volume, the beads were suspended in 20 μL of fresh kinase assay buffer containing 1 μM ATP and 5 μCi of γ^32^P-ATP. Kinase reactions were incubated at 30 °C for 30 minutes with shaking at 800 rpm in an Eppendorf Thermomixer. The reactions were stopped by the addition of 10 μL of 3X Laemmli SDS-PAGE loading buffer and heating at 80 °C for 10 minutes. Twenty-five microliters of each reaction were loaded into a 10 %(v/v) SDS-PAGE gel and proteins were separated for 90-100 minutes at 120 V followed by transfer to PVDF. A storage-phosphor screen was exposed overnight with the PVDF membrane and exposed screens were visualized using a Typhoon imager (GE Lifesciences).

### Mass spectrometric analysis

Samples were prepared and analysed by LC-MS/MS as previously described (Ntoukakis et al., 2009; Piquerez et al., 2014). The mass spectrometry proteomics data have been deposited to the ProteomeXchange Consortium via the PRIDE (Perez-Riverol et al., 2019) partner repository with the dataset identifier PXD025597 and 10.6019/PXD025597.

### Homology modelling and visualization

The homology model of the EFR protein kinase domain (residues 712-1001) was generated using MODELLER (Eswar et al., 2007) implemented in Chimera (v1.15; (Pettersen et al., 2004). The published BAK1 protein kinase domain structures 3UIM (Yan et al., 2012) and 3TL8 (Cheng et al., 2011) were used as templates for the model.

### Software

Figures were prepared using Inkscape (v0.92.3) and GIMP (v2.10.4). Raw immunoblots were converted to TIFF format using BioRad Image Lab (v6.0.1). Plotting and statistical analysis was carried out in GraphPad Prism (v8.3.0 and 9.0.0). Multiple sequence alignments were generated using the ClustalO algorithm in Jalview (v2.10.5). The version of R and emmeans used for analysis of induced resistance data were 4.0.2 and 1.5.3, respectively.

## Author contributions

Experimental work and data analysis: KWB, DC, YK, AM, JS, PD, MB, AP, MFF, BS, VN, LS, AJ, FLHM

Generation of materials: DC, AM, LS

Study design and conception: KWB, YK, AM, BS, VN, AJ, FLHM, CZ

Manuscript writing: KWB, CZ (with comments from all authors)

## Acknowledgements

We thank the TSL Plant Transformation support group for plant transformation, the John Innes Centre Horticultural Services, and Tamaryn Ellick for plant care. All past and current members of the Zipfel group are thanked for fruitful discussions.

## Funding

This project was funded by the Gatsby Charitable Foundation, The Biotechnology and Biological Research Council (BB/P012574/1), the European Research Council under the European Union (EU)’s Horizon 2020 research and innovation programme (grant agreements No 309858, project ‘PHOSPHinnATE’ and No 773153, project ‘IMMUNO-PEPTALK’), the University of Zürich, and the Swiss National Science Foundation grant no. 31003A_182625, and a joint European Research Area Network for Coordinating Action in Plant Sciences (ERA-CAPS) grant (‘SICOPID’) from UK Research and Innovation (BB/S004734/1). Y.K. was supported by fellowships from RIKEN Special Postdoctoral Research Fellowship, Japanese Society for the Promotion of Science Excellent Young Researcher Overseas Visit Program, and the Uehara memorial foundation, and M.B. was supported by the European Union’s Horizon 2020 Research and Innovation Program under Marie Skłodowska-Curie Actions (grant agreement no.703954). B.S. was part of the John Innes Centre/The Sainsbury Laboratory PhD Rotation Program.

---

**Figure S1.**
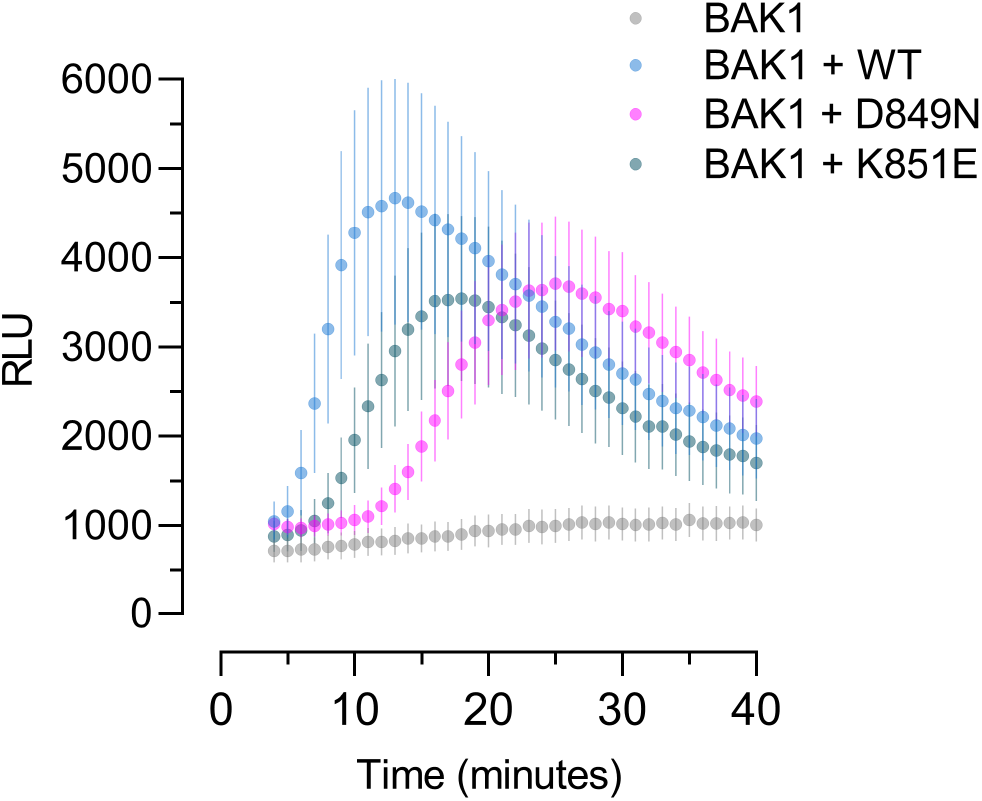
Analysis of the elf18-induced oxidative burst in *N. benthamiana* leaves after transient expression of EFR-GFP or catalytic site mutants. Each EFR variant was co-expressed with Arabidopsis BAK1, and expression of BAK1 alone served as a control for EFR-dependence of the elf8-triggered oxidative burst. Leaf discs were treated with 100 nM elf18 and luminescence was measured for 35 minutes. Points are mean with standard error from six replicate infiltrations.

**Figure S2.**
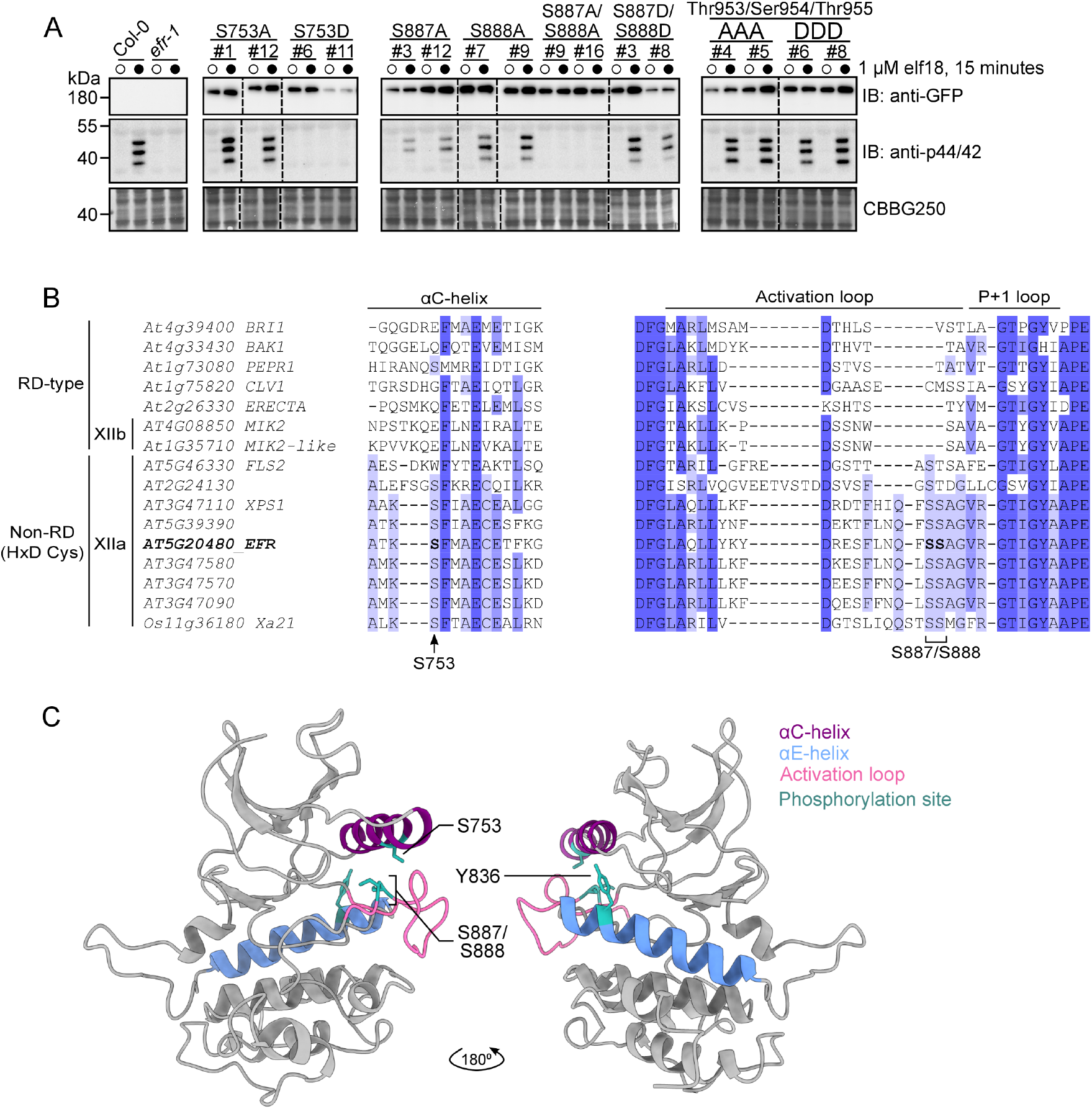
Screen of phosphorylation site mutants for MAPK activation and conservation of regulatory phosphorylation sites. **A**, Immunoblot analysis of MAPK phosphorylation (anti-p44/42) after treatment with mock (open circles) or 1 μM elf18 (closed circles) for 15 minutes in 12-day-old seedlings for non-phosphorylatable (Ala) and phospho-mimic (Asp) mutants of selected EFR phosphorylation sites. Anti-GFP immunoblotting indicates accumulation of the receptor in transgenic plants. Coomassie stained immunoblots are shown as a loading control (CBBG250). **B**, Multiple sequence alignment of Arabidopsis LRR-RKs from subfamily XII with other well-known RKs. Regions of the alignment representing the αC-helix and the activation loop were extracted from an alignment of cytoplasmic domains to reveal conservation of novel regulatory EFR phosphorylation sites. **C**, Homology model of the EFR protein kinase domain showing the location of regulatory phosphorylation sites within important subdomains of the protein kinase.

## Notes

### Competing Interest Statement

The authors have declared no competing interest.

