## Supplemental Table S1 for "ACTIVATION LOOP PHOSPHORYLATION OF A NON-RD RECEPTOR KINASE INITIATES PLANT INNATE IMMUNE SIGNALING"

**Supplementary Table S1.** Representative phosphopeptides identified on immunopurified EFR-GFP**.**

| Phosphosite | Treatment[a] | Mascot ion score | Peptide[b,c] | Observed mass | Actual mass | Charge | Δ ppm |
|---|---|---|---|---|---|---|---|
| Ser683‡ | m/e | 62.46 | K.NNA**pS**DGNPSDSTTLGmFHEK.V | 1109.44 | 2216.87 | 2 | 0.8529 |
| Ser688†‡ | e | 46.68 | K.NNASDGNP**pS**DSTTLGmFHEK.V | 739.96 | 2216.86 | 3 | -2.559 |
| Ser690†‡§ | m/e | 54.29 | K.KNNASDGNPSD**pS**TTLGmFHEK.V | 782.66 | 2344.96 | 3 | 0.2752 |
| Thr691† | m | 58.22 | K.NNASDGNPSDS**pT**TLGmFHEK.V | 1109.44 | 2216.87 | 2 | 1.078 |
| Ser707§ | e | 55.29 | K.VSYEELH**pS**ATSR.F | 729.82 | 1457.62 | 2 | 0.6215 |
| Thr709 | e | 75.46 | K.VSYEELHSA**pT**SR.F | 729.82 | 1457.62 | 2 | -0.6399 |
| Ser753‡ | e | 25.75 | K.HGATK**pS**FmAEcETFK.G | 613.92 | 1838.74 | 3 | 1.316 |
| Ser781§ | e | 31.6 | K.LITVCSSLD**pS**EGNDFR.A | 946.91 | 1891.81 | 2 | -0.52 |
| Ser888‡§ | e | 54.47 | K.YDRESFLNQFS**pS**AGVR.G | 978.44 | 1954.86 | 2 | 1.68 |
| Thr953 | e | 65.51 | K.SILSGc**pT**SSGGSNAIDEGLR.L | 1030.95 | 2059.89 | 2 | 2.563 |
| Ser954‡ | m/e | 84.22 | K.SILSGcT**pS**SGGSNAIDEGLR.L | 1030.95 | 2059.89 | 2 | -1.348 |
| Ser1010† | m/e | 47.6 | K.TTITE**pS**PR.D | 492.72 | 983.43 | 2 | 1.348 |

a, in this study; m, mock treatment; e, 100 nM elf18

b, boldface indicates the phosphorylated Ser or Thr; lowercase 'm' or 'c' indicates oxidized Met and carbamidomethyl Cys, respectively

c, all phosphopeptide spectra were manually inspected to verify site localizations

†, identified in Mergner *et al*. 2020 Arabidopsis phosphoproteome

‡, *in vitro* autophosphorylation site according to Wang *et al*. 2014

§, *in vitro* transphosphorylation by BAK1 according to Wang *et al*. 2014
