## Supplemental Table S3 for "ACTIVATION LOOP PHOSPHORYLATION OF A NON-RD RECEPTOR KINASE INITIATES PLANT INNATE IMMUNE SIGNALING"

Supplementary Table S3. Antibodies used in this study.

| Antibody | Source | Catalog # | Dilution | Incubation time | Reference |
|---|---|---|---|---|---|
| anti-GFP-HRP (clone B-2) | Santa Cruy Biotechnology | sc9996 HRP | 1:2000 | 12-16 hours | n/a |
| anti-p44/42 | Cell Signaling Technology | 9101L | 1:2000 | 2 hours | n/a |
| anti-BAK1 | Custom | n/a | 1:10000 | 12-16 hours | Perraki *et al.* [a] |
| anti-BAK1 pS612 | Custom | n/a | 1:4000 | 2 hours | Perraki *et al.* [a] |
| anti-PR1 | Agrisera | AS10 687 | 1:2500 | 2 hours | n/a |

a, Perraki *et al* . (2018) Nature **561** 248-252
